# GCAT dotplots characterize precisely and imprecisely defined topological features of DNA

**DOI:** 10.1101/2021.07.29.454305

**Authors:** J.I. Collett, S.R. Pearce

## Abstract

Two-dimensional graphical dotplotting is adopted to identify sequence elements and their variants in lengths of DNA of up to 10 kb. Named GCAT for identification of precisely defined short sequences and their variants, its use complements the precise matching of many computational programs, including BLAST. Short reiterated “search” sequences are entered in the Y-axis of the dotplot program to be matched at their identical and near identical (variant) sites in a sequence of interest entered in the X-axis. The result is a barcode-like representation of the identified sequence elements along the X-axis of the dotplot. Alignments of searches and sequence landmarks provide visualization of composition and juxtapositions. The method is described here by example of characterizations of three distinctive sequences available in the annotated *Drosophila melanogaster* reference genome (*www.flybase.org)*: the *Jonah 99C* gene region, the transcript of *Dipeptidase B* and the transposable element *roo*. Surprising observations emerging from these explorations include in-frame STOP codons in the large exonic intron of *Dip*-*B*, high A-content of the replicative strand of *roo* as TE example and similarities of its ORF and the large intron of *Dip*-*B*.

## Introduction

While the currently available computational programs are providing fascinating detail about sequence organization and evolutionary relationships among genomic sequences, these programs largely depend upon precise identities of sequence. But what if sequence configurations are functional with less than precise definition of sequence or if juxtapositions of different combined features provide recognition of functional elements in genomic sequence? Presented here is a graphical method of identifying sequence elements, their variants and their juxtapositions in a topological characterization of sequences of up to about 10 kb. It uses a conventional two dimensional graphical dotplot program to identify particular sequence elements and their variants in their sites within lengths of sequence that often encompass units of function. The resulting dotplot is a linear bar-code-like representation that may be aligned and compared with others of the same or different sequences. Three examples of its use are described here to demonstrate the method and its value in identifying sequence elements and visualizing their relationships.

The method co-opts the Y-axis of a two-dimensional dotplot program for use as a purpose-designed “search sequence” to identify particular sequence elements within a sequence of interest entered in the X-axis. It is thus dubbed “GCAT”. The sequences searched here as examples of the ‘GCAT’ analysis, were chosen from the annotated genome of *Drosophila melanogaster* (*www.flybase.org*). Each analysis follows directly from the detail of their annotations, from conventional dotplotting, from database identifications of their functional elements and from various BLAST (Basic Linear Alignment Sequence Tool) searches. The interests of each analysis are to relate sequence structure to function and to understand their evolutionary histories.

Three distinctively different genomic sequences were chosen to describe possible applications of GCAT visualization and the contributions it offers in characterizing sequence topology in identifying sequence configurations that may elude the precise requirements of computational identification. Characterization of the 99C region of the *Jonah 99C* genes (*99Ci, Cii* and *Ciii)* describes the topological context of these regulated members of the large dispersed gene family (Carlson and Hogness,1985) and offers example of identifying likely footprints of historic sequence amplification. An exploration of the transcript sequence of the constitutive gene *Dipeptidase-B* focuses upon the distinctive nucleotide composition of its introns and the possible roles of its large exonic intron in maintaining dynamic stability in transcription and translation. This then provides a basis for comparing and contrasting this intron sequence with the apparently active and widely dispersed transposable element (TE) *roo*, described in some detail by Kaminiker *et al.* (2002). In encompassing domains of genetic function, each analysis, aligned with their annotated landmarks, offers visualization of their topological contexts and of functional elements that may also reflect their evolutionary histories.

## Methods

### Modified use of Conventional Dotplotting in GCAT analysis

The Gilbert (1989) DottyPlot program, commonly used to visualize sequence alignments in two-dimensional graphical matrices, is adopted here to identify short sequence elements within sequences of up to about 10 kb. A brief description of the conventional use of dotplots may be helpful in understanding how the dotplot program also lends itself to identifying short sequence configurations and their sites in these GCAT analyses.

Conventional dotplotting is most often used to identify commonality within and between sequences. Dotplot programs match the base composition of one sequence with itself or with another entered in the X- and Y-axes. It matches the base composition of the sequences in the X- and Y-axes within defined windows that progress base-by-base along each axis. Identity of a match is defined by the specified number of matching bases in each match window. Identity in the Gilbert (1989) program also requires the same order of bases within matching windows, but the order may be interrupted anywhere along its length within a window when the match requirement is set to be less than perfect. Windows in which the preset identity requirement is met are represented by dots at the X- and Y-coordinates in the resulting two- dimensional graph. Complete identity of the sequences entered in the X- and Y-axes are then indicated by a straight diagonal line through the space of the coordinates while reiterated sequence and direct repeats appear as patterns and subsidiary offset lines that indicate their positions and lengths within both the X and Y sequences. An example of such a conventional dotplot may be seen in Figure 1.

**Figure 1.**
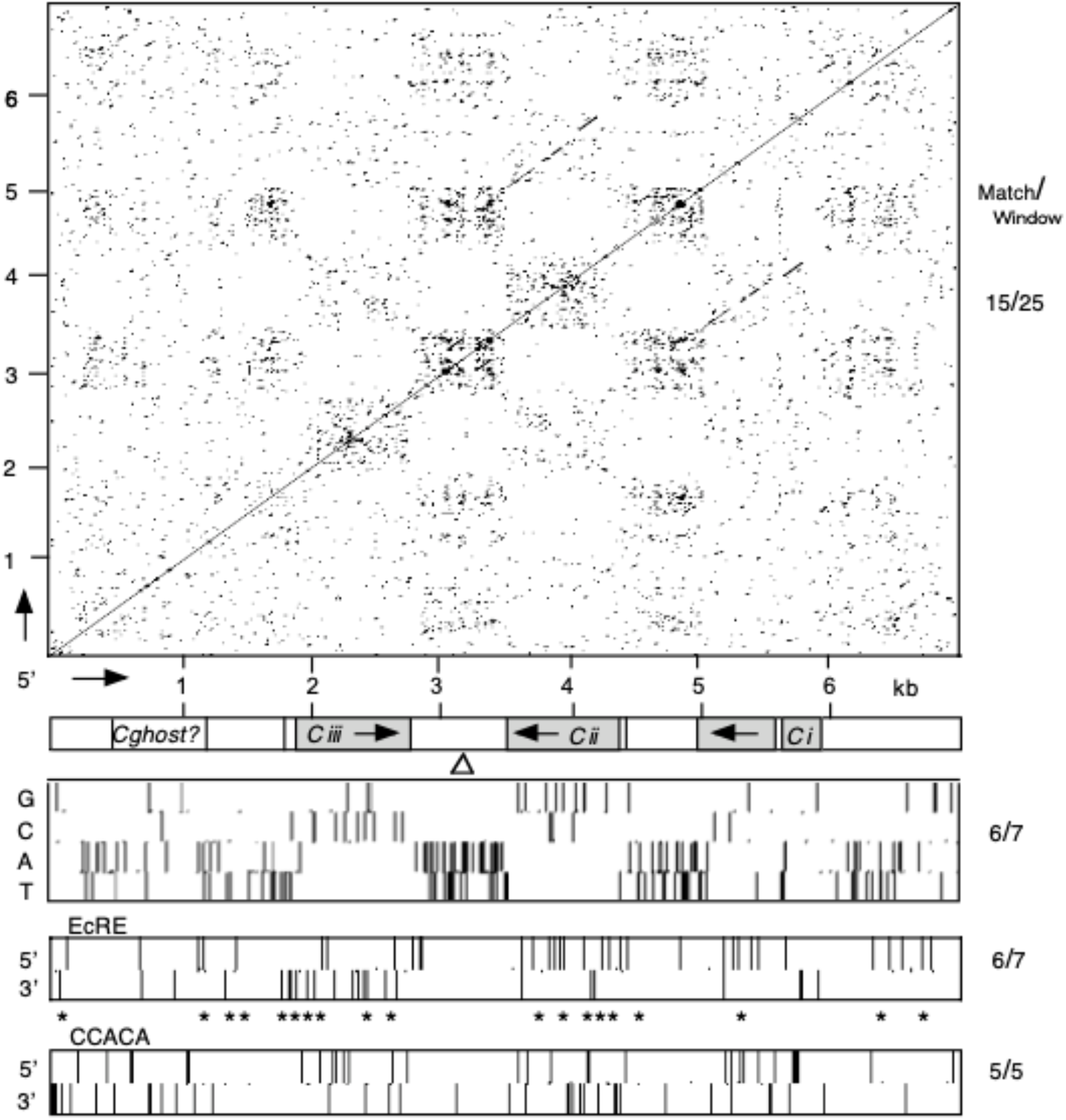
Conventional 2D dotplot of the *Jonah 99C* gene region of *D*. *melanogaster* and its GCAT explorations. The 99C chromosomal region as analyzed here extends 6,962 bp from the 3’UTR of the gene *trp* to the 5’UTR of the gene CG17722. (*www.flybase.org*). Top panel, a two-dimensional (2D) dotplot of the *99C* sequence entered in both the X and Y axes in the orientations indicated, requiring a match of 15 bases in successive 25 base windows. Each match is indicted by a dot. The panel aligned below the X-axis is a map of its sequence landmarks including the coding sequences of the *Jonah* genes, *Ci*, C*ii* and *Ciii* (grey) the directions of their transcription (arrows), the 5’UTRs of Cii and *Ciii*, a 57 bp intron in *Ci*, an inverted triangle indicating a 12 bp palindrome, and a boxed *Cghost?* (as described in text). **GCAT** panel. A search of sequence composition, using a four-part sequence of GGGC, CCCG, AAAT, TTTA, each reiterated in tandem, and entered in the Y-axis with a match requirement of 6 bases in 7 base windows, aligned with the landmark map above. **EcRE** panel. A search for ecdysone response element-like sequences (EcRE) using a reiterated sequence of 7-base ECRE, GAGGTCA (5’) and its reverse complement (3’) with a match requirement of 6 bases in 7 base windows, aligned with landmark map above. Stars (*) indicate sites of perfect EcREs or single base variants. **CACCA** panel. A search for 5’ and 3’ orientations of the sequence CACCA that characterizes the coding sequence of *Jonah 99Ci* with a match requirement of 5 bases in 5 base windows, aligned with the landmark map above.

In these GCAT analyses several differences in the use of dotplot programs are introduced in order to identify short sequence elements and their positions within a DNA sequence of interest. The sequence to be searched is entered in the X-axis of the dotplot program, while the “search” sequences are entered in the Y-axis. But in order to see the positions of these identified search sequences, reiterated copies are entered as the search sequence to result in a series of dots appearing as a vertical line in the Y- axis above its position in the “searched” sequence in the X-axis. The match requirement may be set to identify only perfect matches of the search sequence or to allow “wobble” in identifying similar but less than perfect identities of the search sequence in the X-axis sequence.

Further, several searches of the X-axis sequence may take place simultaneously by expanding the Y-axis to include more than one search sequence, allowing graphical identification of two or more sequence identities and their juxtapositions. The resulting dotplot looks like a barcode, or in the case of several simultaneous searches, as a stacked barcode. (It may also be noted that in these stacked barcodes, the partial sequences of the transitions between one search sequence and another on the Y-axis may identify overlapping search sequences appearing as isolated apostrophes in the stacked interfaces, as in Figure 1c).

These GCAT searches may also be aligned with other information about the X-axis sequence, as shown here in alignments with their mapped sequence annotations (*www.flybase.org*).

### *Drosophila melanogaster* sequences of GCAT analyses. *Jonah 99C* gene region

(Figure 1). This 6,966 bp sequence (3R: 29,920,289 - 29,927,255) extends from the 3’UTR of the gene *trp* to the 5’UTR of CG15522 (http://flybase.org, R6.27, 2019-02). This is the orientation of its GCAT searches, and includes the genes *Jonah 99Ci* (FBgn0003358), *Jonah 99Cii* (FBgn0003356) and *Jonah 99Ciii* (FBgn0003357), as annotated. The search sequences and parameters are provided in the legend of Figure 1.

### Transcript of *Dipeptidase-B*

(Figure 2). This 3,389 nt transcript of the annotated gene *Dip*-*B* (FBgn0000454) located on chromosome 3R:13,782,288 - 13,785.676 (*www.flybase.org*, FB2019-02, R6.27) was searched in its 5’orientation. The search sequences and match parameters are described in the legend of Figure 2.

**Figure 2.**
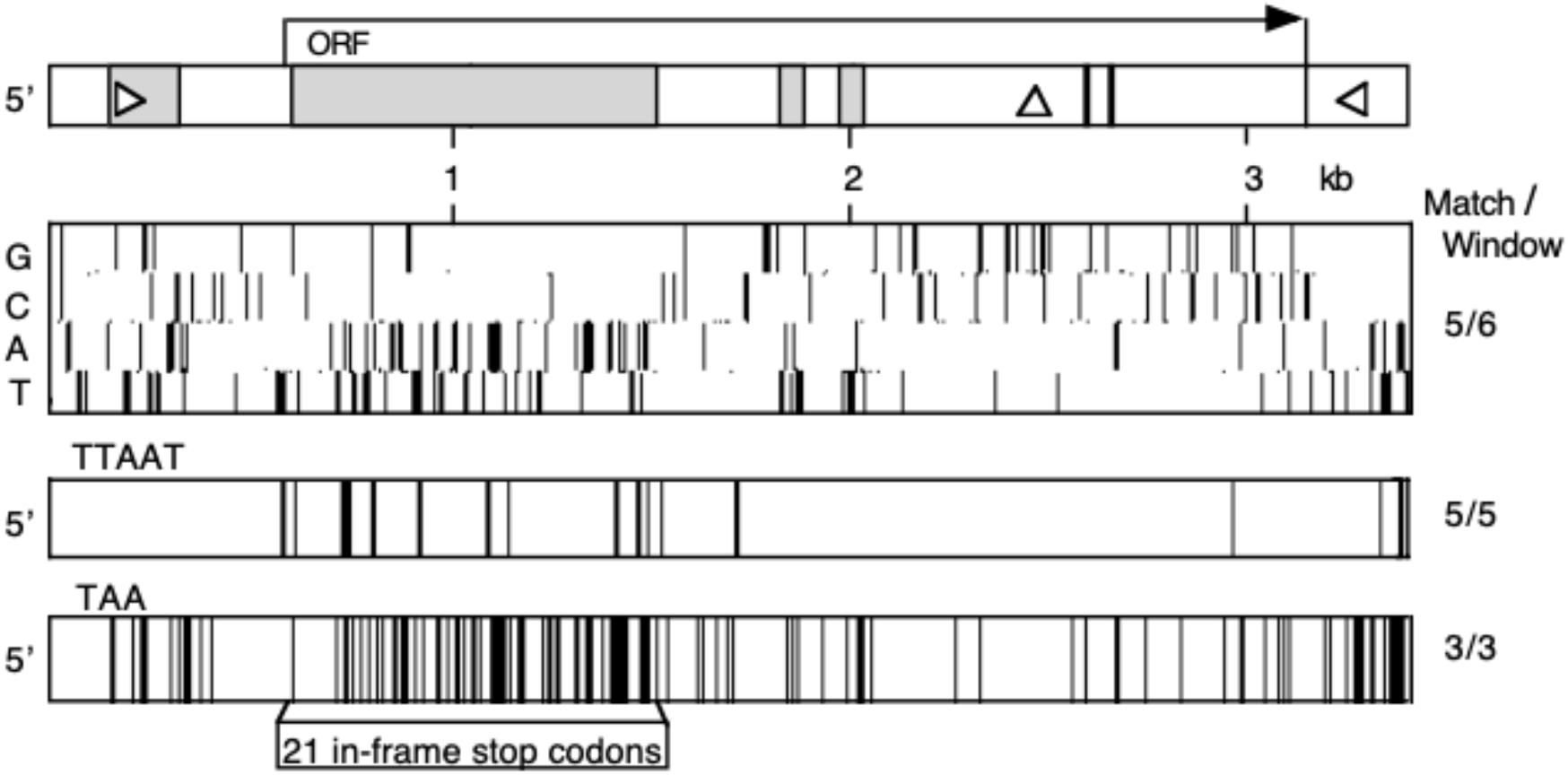
GCAT characterization of the *Dipeptidase-B* transcript of *D*. *melanogaster*. (Top panel) Sequence map of 3,390 bp transcript including its five exons (white), four introns (grey) and 1.5 kb ORF (http://flybase.org), a pair of 12 nt inverted repeats in 5’ and 3’ UTRs (triangles), a 12 nt palindrome (vertical triangle) and putative active site of coding region (http://prosite.expasy.org) within thick vertical lines. This sequence was entered in the X-axis of each of the following GCAT searches. The results were aligned with this map. **GCAT** panel. Result of search for the sequences GGGC, CCCG, AAAT and TTTA, each reiterated in tandem and positioned at the level of the Y-axis indicated by G, C, A, T. Matches required identity of 5 bases in 6 base windows. **TTAAT** panel. Result of search for the intron excision signal sequence, TTAAT, tandemly reiterated in the Y-axis. Matches required identity of 5 bases in 6 base windows. **TAA** panel. Result of search for the STOP codon TAA, tandemly reiterated in the Y-axis.

### LTR-transposon *roo{419}*

(Figure 3). The apparently full-length 9078 nt LTR-transposon *roo {419}* (Kaminiker *et al.*, 2002), identified in chromosome 2L:19,703,592 - 19,712,669 as FBti 0019205 (*www.flybase.org)* in the 5’ orientation of its ORF was used in the searches presented in Figure 3 as described in its legend.

**Figure 3.**
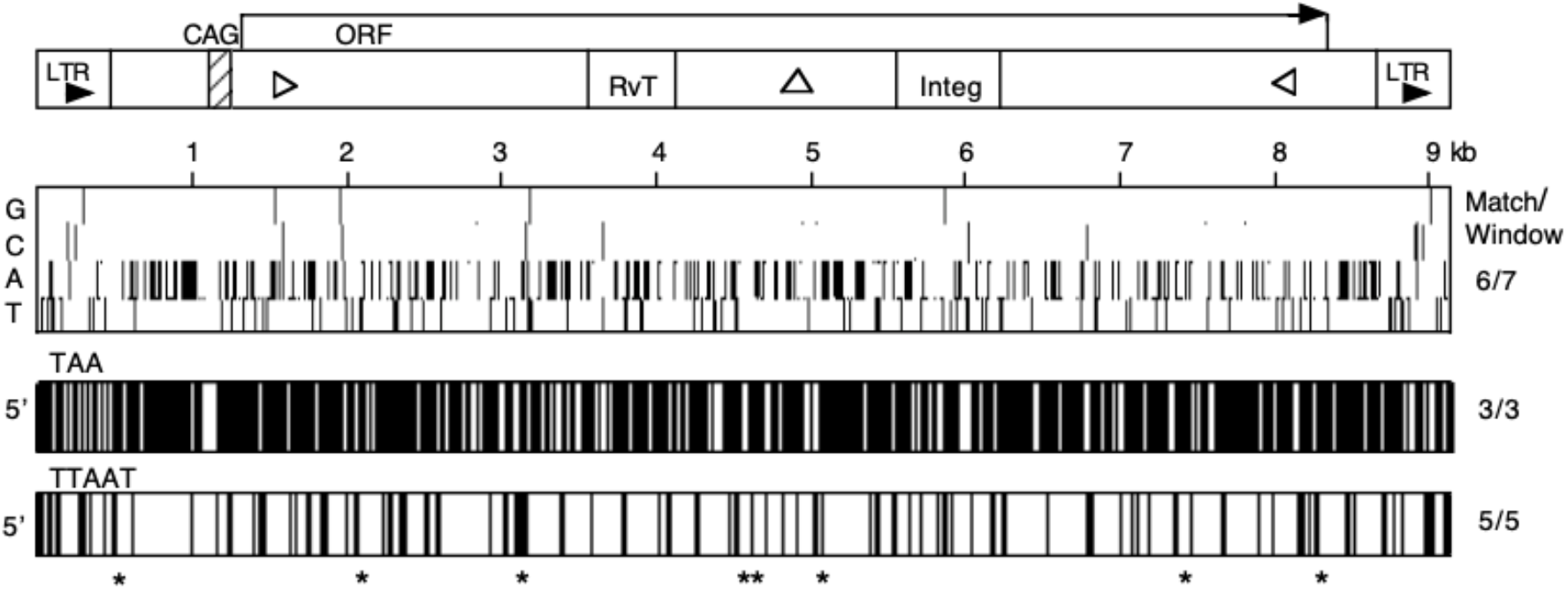
GCAT Characterizations of Transposable Element *roo* of *D*. *melanogaster*. (Top panel) Map of features of the 9078 nt sequence of the transposable element *roo{}419* including Long Terminal Repeats (LTR), a CAG-rich sequence and its single 7.2 kb ORF. Sites identified within ORF are sites of putative coding identified as reverse transcriptase (RvT) and integrase (Integ) (*InterPro* databases, www.ebi.ac.uk), a pair of 20 nt inverted repeats (triangles) and a 14 nt palindrome (vertical triangle). Aligned below map are GCAT characterizations of the *roo* sequence. **GCAT** panel. Result of search for sequences GGGGC, CCCCG, AAAAT and TTTTA, each reiterated in tandem and entered in the Y-axis at the levels indicated by G,C,A,T. Matches required identity of 6 bases in 7 base windows. **TAA** panel. Result of search with tandemly reiterated STOP codon TAA entered in the Y-axis. Matches required identity of 3 bases in 3 base windows. **TTAAT** panel. Result of search for tandemly reiterated intron excision signal TTAAT in the Y-axis. Matches required identity of 5 bases in 5 base windows. * indicates sites of TTAAT configurations followed by a substantial polypyrimidine tract upstream of an AG.

### BLAST and database searches and GCAT graphic representation

Sequences were submitted to the NCBI BLAST (bl2seq) basic BLAST program (http://blast.ncbi.nlm.nih.gov) in lengths ranging from 500 nt to the full lengths to identify repeats in both orientations using both standard and low stringency match parameters. Dipeptidase-B was identified as a member of the M17 peptidase family in the MEROPS database (*http://merops.sanger,ac,uk)* (Rawlings and Barrett, 1999). Its active site is indicated in Figure 2. The putative functional coding regions of the ORF of *roo* were identified in an InterPro scan of protein databases (*www.ebi.ac.uk*). GCAT dotplots were generated using the Gilbert (1989) Dottyplot program for Mac computers (OS 9) and transferred in SimpleText to Appleworks 6.0 for scaling and labeling.

## Results

### GCAT Exploration of the *Jonah 99C* gene region

The *Jonah* genes of *Drosophila* comprise a dispersed family of genes active in the gut (Carlson and Hogness,1985) that code trypsin-like proteases (www.Flybase.org). Those located in the 99C region were shown to be nearly exclusively transcribed in the posterior mid-gut (Akam and Carlson, 1985) and the correspondence of these *Jonah* genes in size, activities, sites of tissue localization and map location (Beckman and Johnson, 1964; Hall, 1988a) to a complex of related Leucine aminopeptidases (LAP) (Seymour-Jones and Collett, 1992) compellingly suggest that these *Jonah* genes (and/or those in 99F) code the multi polymeric LAP. Extensive physiological analysis of LAP (Hall, 1988b; Collett, unpublished) also demonstrated LAP activity to be differentially regulated between the sexes of adults by ecdysone in females. Responsive regulatory sequence elements as well as clues about the region’s ancestral history are therefore likely to be embedded within the sequence of the *Jonah 99C* region.

Exploration of the *Jonah 99C* region began with a conventional dotplot and extensive BLAST alignments of sequences within the region to establish a starting point for the particular sequence searches with GCAT analysis. This conventional two-dimensional dotplot may be seen in the top panel of Figure 1 together with, below, the aligned map of the annotated genes of *99C*. In this dotplot the region’s sequence is matched to itself in the same orientation using a permissive match requirement of 15 bases in 25 base windows. The diagonal line through the graph represents the match of the sequence with itself in the X- and Y-coordinates, while the similarities of *Ci* and *Cii* are indicated by the offset diagonals above and below each X-axis sequence at their positions in the Y-axis. It should also be noted that since genes *Cii* and *Ciii* are present in the opposite orientation, their identities are not indicated in this dotplot.

Thus the first GCAT analysis provides a simple characterization of the nucleotide composition of the region. The search sequence of GGGC, CCCG, AAAT and TTTA, each reiterated in tandem, was entered in the Y-axis of the dotplot program. The choice of requirement for identity matching followed from visual inspection of the commonly occurring sequence configurations in the region and was set to identify (at least) 6 bases in windows of 7 bases. This Identification therefore required, for example, three adjacent Gs, a C, and two more Gs in any order within each base window. The resulting barcode-like representation of the distributions of each nucleotide concentration may be seen in the “GCAT” panel of Figure 1.

The alignment of the base composition of the 99C region with the coding and intergenic sequences of the region demonstrates the distinctively different composition of each. Intergenic sequence is clearly AT-rich, while the G and C barcoding of the three coding sequences describes the distinctive patterns of each gene. Here, the complementary patterns of *Cii* and *Ciii* indicate both their near identity and their inverted orientations while also indicating many differences with the *Ci* gene. The compositional GCAT representation in its alignment with the 2D dotplot above also describes the composition of its checkerboard pattern. It arises from AT-dense intergenic sequence bounding the GC-rich gene sites of the region.

Thus the first GCAT analysis provides a simple characterization of the nucleotide composition of the region. The search sequence of GGGC, CCCG, AAAT and TTTA, each reiterated in tandem, was entered in the Y-axis of the dotplot program. The choice of requirement for identity matching followed from visual inspection of the commonly occurring sequence configurations in the region and was set to identify (at least) 6 bases in windows of 7 bases. This Identification therefore required, for example, three adjacent Gs, a C, and two more Gs in any order within each base window. The resulting barcode-like representation of the distributions of each nucleotide concentration may be seen in the “GCAT” panel of Figure 1.

The alignment of the base composition of the 99C region with its coding and intergenic sequences demonstrates the distinctively different composition of each. Intergenic sequence is clearly AT-rich, while the G and C barcoding of the three coding sequences describes the distinctive patterns of each gene. Here, the complementary patterns of *Cii* and *Ciii* indicate both their near identity and their inverted orientations while also indicating many differences with the *Ci* gene. The compositional GCAT representation in its alignment with the 2D dotplot above also describes the composition of its checkerboard pattern. It arises from AT-dense intergenic sequence bounding the GC-rich gene sites of the region.

These GCAT-defined patterns of nucleotide composition led to BLAST searches to establish their precise sequence identities. The coding of genes *Cii* and *Ciii* are 99.3% identical in their inverted orientations while differing substantially in their short UTRs. Their conspicuous differences with the coding sequence of *Ci* reflect a 50% difference in coding as well as its short mid-sequence AT-rich intron.

Further BLAST analysis of the region’s sequence in overlapping 500 bp lengths found no other substantial identities throughout the region even with low stringency match parameters. But BLAST searches did identify a 12 bp palindrome at the center of the region mid-way between genes *Cii* and *Ciii*. Its position (see Figure 1) suggests it to be the site of insertion of one end of the apparent inverted duplication of the *Jonah Cii* or *Ciii* gene. This possibility is returned to below.

How useful GCAT searches might be in identifying the short sequence configurations associated with their interactions with, for instance, protein regulators, was taken up with a search for the 7 nt sequence identified by Vogtli *et al.* (1998) to be an ecdysone response element in *D*. *melanogaster*. They showed that the core sequence motif, GGAGTGA, as well as some of its single nt variants responded to ecdysone in their *Drosophila* assay system. Thus the possibility that this sequence and/or its variants might be involved in regulating expression of one or more of these *Jonah* genes as is the case of LAP (Hall,1988b) and in adult female guts (Collett, unpublished) was taken up using their defined 7 nt ecdysone response element (EcRE) as a search sequence. This 7-nt sequence was entered in the Y-axis as a two-part search sequence, consisting of its 5’ orientation and its 3’ reverse complement, with a match requirement of 6 bases in 7 base windows. This search thus identified both the exact sequence and its single nt variants in each orientation (Figure 1, EcRE panel).

Of the 53 EcRE-like sites identified in this search, 19 sites are the sequence itself or its single nt variant, as indicated. The sites are clearly concentrated in and near the active genes *Cii* and *Ciii* in both orientations and are nearly absent from *Jonah Ci* and its closely associated non-coding sequence. These sites, including particularly the 5’ upstream sites of *Ciii* thus confirm the possibility of the *Cii* and/or *Ciii* genes’ ecdysone responsiveness, as well as indicating how the three *99C* genes may be differentially regulated in adult males and females.

The distribution of these EcRE sites within the coding sequences of *Ciii* and *Cii* also suggests another intriguing possibility. Could these intra-coding sites draw in and concentrate EcRE complexes, thus increasing their effectiveness in associating with the response elements in positions that promote greater transcriptional activity? Identification of this unexpected possible dual functionality of coding sequence may be a valuable observation to have emerged from this visualization.

Use of GCAT analysis to identify sequence patterns that might reflect their history was then taken up in a search for sequence upstream of *99Ciii* that particularly characterizes sequence of *Jonah Ci* as, perhaps, vestiges of its sequence included in the distal end of region’s historic duplication. Using a sequence configuration found to be conspicuous in each of the three *Jonah* genes, CCACA, but largely absent from intergenic *99C* sequence, the CCACA panel of Figure 1 shows that indeed the 5’ distal region of *99C* includes this CCACA configuration in both orientations similar to their occurrence in the currently active *Jonah 99Ci* gene. This identification therefore confirms the possibility that at least some of the *Ci* gene was present in the originating duplication of this *Jonah* gene region.

However, the more persuasive evidence of the inclusion of some or all of the *99Ci* gene in the region’s inverted duplication may be seen in the coincidence of these CCACA configurations with the more gene-like sequences identified in each of its GCAT characterizations. Looking up through the search for EcREs to the GCAT characterization of nucleotide composition and then to the 2D checkerboard of the region, it will be seen that the patterns of each correspond more to gene-like sequence than to intergenic sequence in this 5’ part of the *99C* region. Together, these coincident patterns offer strong suggestion of the distal end of the historic event that established this region. This suggestion of now disintegrated identity is therefore noted on the map of the region’s annotations as “*Cghost?”* as perhaps the site of the birth and death of a once active gene.

### GCAT Exploration of the *Dipeptidase B*

*Dipeptidase-B*(*Dip-B)* codes one of a cohort of ‘Dipeptidases’ that service the final step of protein catabolism in flies (Collett, 1989). The apparent ubiquity of transcript (http://flybase.org; Graveley *et. al.*, 2011) and of measured enzymatic activity in stage and tissue of the fly (Laurie-Ahlberg, 1982; Hall, 1988a) characterize the gene as constitutive. Together as a cohort of Dipeptidases, their intracellular activites in hydrolyzing small peptides to free amino acids are regulated in several ways that maintain amino acid in small peptides as an osmotically tolerable reservoir in the fly (Collett,1989). Dip-B hydrolyses a variety of peptides at rates limited by the amounts of enzyme present. The particular interest here was therefore to explore whether alternative transcript processing might provide alternative forms and activities of Dip-B, for instance, in any of the fly’s differing requirements, as through development, or in its more extreme environments as in overwintering (Collett and Jarman, 2007). Thus, GCAT characterization of the sequence structure of *Dip*-*B’s* transcript with special interest in its 4 introns might reveal capacity to provide alternative transcripts and activities.

Landmarks of the *Dip*-*B* transcript (www.flybase.org) are presented in Figure 2. In common with many *Drosophila* genes, *Dip*-*B’s* annotation includes a substantial intron in its 5’UTR and another nearly one kb intron immediately downstream of the 5’ initiating ATG of its ORF and two short introns mid-way along its coding sequence. These together account for about 1.5 kb of its 3.4 kb transcript. The 317 identified cDNAs (www.flybase.org) of *Dip*-*B* correspond to the commonly processed transcripts most probably in flies reared in standard laboratory conditions. They differ only in their differently excised 5’UTR intron. BLAST searches of the transcript sequence reveal a palindrome near the middle of the transcript (and mRNA) and a pair of inverted repeats present in the transcript’s 5’UTR intron and in its 3’UTR. The entire transcript is the subject of these GCAT searches.

Like the GCAT exploration of the *Jonah 99C* region, these began with a characterization of the transcript’s sequence topology. A 4-part search sequence (Figure 2, legend) was used to establish the distribution of nucleotides throughout its length and to identify sites of concentration of each base. Its alignment with the landmark map of the transcript’s annotations indicates striking GC richness in its coding in contrast to the AT density in its non-coding sequences. These differences are reflected in the distinctive ‘barcoding’ of its coding and non-coding sequences.

The first exploration of *Dip-*B was to discover whether the sequence of the large 5’ exonic intron could allow alternative splicing and therefore provide alternative coding and alternative activity or regulation of Dip-B, as in times of environmental stress and overwintering. Following from its AT composition (64.5%), a search for sequence that might allow alternative excision was carried out. Using the commonly found excision signal sequence TTAAT, as a search sequence with a match requirement of 5 bases (in any order) in 5 base windows, a number of sites were identified in the transcript’s large intron (see TTAAT panel, Figure 2).

These sites led to a visual search for the exact TTAAT configurations that are also followed by the commonly found polypyrimidine tract preceding an AG excision site that re-establishes the IN-frame coding of the ORF’s originating ATG. Two of these configurations near the 3’ end of the intron’s sequence fit these criteria. But, while both would re-establish the IN-frame coding of the ATG originating ORF, the first and more 5’ of these includes an IN-frame STOP codon immediately following its would-be 3’ AG excision site! Thus a definitive answer followed from this GCAT search. The sequence structure of this long intron can only deliver the coding and the activity of *Dip*-*B*, as annotated.

The three shorter introns of the *Dip*-*B* transcript are also of interest. The two short AT-rich introns in the middle of the transcript’s coding (see Figure 2) also contain STOP codons that would be IN-frame were their excision to fail, thus stopping translation, in these cases, before inclusion of *Dip*-B’s active site. But the intron of the 5’UTR is alternatively spliced, as indicated by the identification of *Dip*-B’s RNAs in the fly’s transcriptomes (Graveley *et al.*, 2011). Although these RNAs were probably sampled from flies raised in standard laboratory conditions, their differences indicate the possibility that indeed different 5’UTRs may affect transcription rates, transcript stabilities and/or transcript turnover rates, thus affecting the amount of active Dip-B in the fly, as in response to the range of environmental circumstances the fly encounters in its natural environments. That responsiveness might well contribute to Dip-B’s role in releasing essential amino acid in the fly.

While these discoveries offer interesting detail about how introns may both facilitate transcript function and prevent its mal-function, they offer little solution to the puzzle of the sizes and positioning of the common 5’exonic introns, characterized by *Dip*-*B’s* large intron. Pursuing the distribution of its AT composition further, a search for the STOP codon TAA was taken up (see TAA panel, Fig. 2). In identifying ATA, AAT and TAA triplets throughout the transcript, the result indicates their varied distribution throughout the transcript. This indicates how T and A are distributed throughout the transcript and most particularly are concentrated in introns and the UTRs. Most interestingly, however, is that a further visual count of all STOP codons (TAA, TGA and TAG) in the long intron indicated that among very many, a striking 21 stop codons are IN-frame with the originating ATG of the coding sequence. Thus clearly, if, or perhaps when, excision of this intron fails, so too does its translation.

A small survey of some of the many *D*. *melanogaster* genes carrying these long 5’ exonic introns indicated that indeed their IN-frame STOP codons appear to describe a general rule about the composition of these long 5’ exonic introns. This then raises the further curiosity: how did introns get there and what does their baggage offer the fly? In *Dip*-*B’s* case, for example, about one-third of the transcript goes nowhere but is returned to cellular space. This led to comparing the sequence structure of these long introns to the sequence structure of transposable elements.

### Exploration of the Sequence Topology of the TE *roo*

Occupying in aggregate about 20% of the *D*. *melanogaster* genome (Barron *et al.*, 2014), questions about how Transposable Elements (TEs) may affect and contribute to genes and genomes and their evolutionary dynamics through at least their insect ancestry were then taken up in further exploring the usefulness of GCAT methodology. The well-studied LTR retrotransposon *roo* (Kaminniker *et al.* 2002) was chosen for this analysis, using as its example *roo{419}*, present in the reference sequence of *D*. *melanogaster* in a long intron of the gene CG46244, a transcriptionally active gene of unknown function (*www.flybase.org*). Differing in its sequence by only a few nucleotides, Ashburner and Bergman (2009) chose this element to represent *roo* as its canonical sequence in their compilation of TEs in this *Drosophila* species.

The exploration began with the identification of sequence landmarks in its 9 kb (Figure 3). These include identical (direct) Lateral Terminal Repeats (LTRs) and a single 7 kb ORF that in its mid-region includes coding identified in an InterPro scan (www.ebi.ac.uk) as reverse transcriptase and integrase. The ORF is also claimed (Frame *et al.*, 2001) to carry coding for an envelope protein near its 3’ end. Other intriguing details of its sequence, identified in BLAST searches, are inverted repeats near the 5’ and 3’ ends of its ORF and a 120 nt CAG-rich sequence immediately upstream of the ORF’s originating ATG. Its LTRs include a number of short repeats and several longer direct repeats lie between the 5’ and 3’ ends of the ORF and the LTRs. Not included in this sequence map (Figure 3) are sites matching RNA sequences within the ORF, none of which coincide with the ORF’s originating 5’ ATG (http://flybase.org). These elements clearly suggest contextual sequence complexity in the ORF. This is the particular subject of this exploration.

Following from the GCAT characterizations of the *Jonah* gene *99C* region and *Dip*-*B’s* transcript, the surprise of this characterization of *roo* was in the nucleotide composition of its ORF. Using a more stringent search requirement than for the gene sites of *Jonah 99C* and *Dip*-*B* in defining its sequence topology, this GCAT search required matches of 6 bases in 7 base windows of reiterated GGGGC, CCCCG, AAAAT and TTTTA. A number of features stand out in this characterization (GCAT panel, Fig. 3). A is abundantly conspicuous throughout most of the element, while its LTRs look more gene-like in their nucleotide composition. Its ORF is 68.5% A and T and includes few of the concentrations of G and C that characterize genomic genes. A mid-region palindrome and inverted repeats in the 5’ and 3’ ends of the ORF are also intriguing topological details. Particularly notable among these details is that *roo*, among many of the longer TEs of *Drosophila*, carries a predominance of A in its ORF (and replicative strand) (Ashburner and Bergman, 2009). This density of A is thus not an exception, but characterizes many TEs.

It should also be noted that the sequence composition of the *roo* ORF is strikingly similar to *Dip*-*B’s* long 5’exonic intron. This led to detailed comparisons of their T and A configurations. The first search, for TAA STOP codons, used the same search parameters used to identify *Dip*-*B’s* STOP codons. And like *Dip*-*B’s* large intron, the *roo* ORF is densely packed with TAA configurations (Fig.3). But perhaps more striking is that not one of these STOP codons in its 7kb, is IN-frame with the originating ATG of this ORF! A visual count of all its STOP codons, TAG, TGA and TAA, number about 65 per kb throughout its length. Perhaps more remarkable about this number is the vulnerability they bring to the ORF coding in the occurrence of single and doublet indel mutation.

This observation thus led to question whether and how this ORF of *roo* may differ from genomic ORFs. A search for sequence that might allow excision of perhaps functional sequences from within the ORF was therefore undertaken, using the search protocol to identify the TTAAT excision signal sequences of *Dip*-*B’s* long intron. The resulting barcode pattern (TTAAT panel, Fig 3) prompted further visual searches for the commonly found coincident polypyrimidine tracts and in-frame AG excision sites of long genomic introns. These coincident configurations were indeed found, as indicated in Figure 3. Moreover, several of their sites encompass the apparent coding sequences of reverse transcriptase and of integrase. Is it, then, the case that from these readily transcribed ORF sequences, lengths of transcript coding the essential activities of TEs are released from the 7 kb ORF sequence for independent translation and activity in their nuclei? Some evidence of this may be found in the sampling of RNA from early embryos that match not the originating 5’ sequences of the ORF, but lengths of the ORF’s mid-region of *roo{419}* itself in an intron of gene CG46244 (*www.flybase.org*).

Clearly, the sequence structures of *Dip*-*B*’s long intron and the *roo* ORF and their configurational organization are clearly strikingly similar. This coincidence does indeed suggest that invading TEs took up compatible residence within genes forming TEs. But do these TE sequences now provide, in their similar composition, a role similar to introns in the dynamics of nuclear activity? A clue may be in the further similarity of the lengths of the two sequences. In *Dip*-*B’s* case, about one-third of the transcript is disposed of in intron excision before its messenger maturation, while in *roo*, it seems likely that a large part of its ORF is also disposed of in the maturation of its functional transcript. Both *Dip*-*B’s* large intron and *roo*’s ORF seem likely to be osmotically safe dynamic reservoirs of nucleotide substrate, on site and accessible for incorporation in nucleic acid synthesis. This possibility makes sense of what otherwise seems like chromosomal baggage.

## Discussion

The principal interest of this study was to develop a way to identify sequence configurations and their variants and to visualize the positional relationships of these configurations to complement computational analysis in understanding relationships of structure and function. For this we adopted the two-dimensional graphical dotplot program (Gilbert,1989) created to compare one sequence with itself or with another in its X and Y axes. We used the Y-axis for short defined and reiterated sequences to search for their similar or identical sequences in a sequence of interest entered in the X-axis. These defined, and thus dubbed “GCAT” sequence searches, provide a linear barcode-like representation of the X-axis sequence that may be aligned with other searches and maps of landmarks of the sequence to show positions and juxtapositions.

To test the usefulness of this method of exploring and visualizing sequence, we took up three distinctively different and long-standing curiosities in *Drosophila* genetics. The first was an exploration of the *Jonah 99C* gene region, one of the several dispersed gene sites of the *Jonah* gene family that are active in the fly’s gut (Carlson and Hogness,1984). Following this a characterization of the transcript sequence of the constitutively active *Dip*-*B* was taken up with particular interest in sequence structures of its introns including its long 5’ exonic intron. This then led on to examining its resemblance to a currently active transposon, *roo*, as example, perhaps, of its ancestral origin. The discoveries emerging from these explorations persuaded us of the value of these visualizations of sequence, their configurations, their variants and their positional relationships in prompting speculation about relationships of sequence structure and genomic function. These discoveries and their possible significance are taken up here.

The dispersal of the *Jonah* genes of *D*. *melanogaster* has undoubtedly taken place through a long insect ancestry, but its currently active gene members in the 7 kb of the *99C* chromosomal region, *99Ci, 99Cii* and *99Ciii*, probably reflect relatively recent events. Further, the matching profile of these *Jonah* genes to the Leucine amino peptidases (LAP) (Hall,1988a), as their likely gene sites, suggested opportunity for exploring the region’s sequence structure using the search capacity of GCAT analysis to identify possible sites of LAP’s ecdysone responsiveness in adult females (Hall, 1988b) and to track the region’s recent history.

A striking checkerboard pattern of the region in a conventional 2D dotplot provided a tantalizing characterization of its topology and led to searches for its particular detail (Figure 1). First, a GCAT representation of the region’s nucleotide composition aligned with a map of the annotations of the region (*www.flybase.org*) demonstrated clear distinction of the GC-rich coding sequence and the AT-rich intergenic sequence of the 2D checkerboard pattern of the region. This barcoding also demonstrates GCAT’s simple graphical distinction between the three coding sequences. The near identity of *Jonah Cii* and *Ciii* but within an apparent inversion and their differences with *Jonah Ci* are immediately apparent. Together they also define some of the checkerboard.

This characterization of the region also provided a topological map for further exploration of the region. Its first search was for the 7 nt sequence identified by Vogtli *et al.* (1998) found to be effective together with some of its single nt variants as ecdysone response elements (EcRE). In a search to identify both the perfect sequence and its single nucleotide variants, a number of sites with a perfect or a single variant match were found upstream of the *Ciii* and, interestingly, within *Ciii* and *Cii* while nearly absent from *Ci*. The distribution of these sites may therefore explain the observed differences in ecdysone-responsive LAP activities between the sexes of adults.

But a further intriguing detail emerging from this GCAT search is the occurrence of EcRE-like configurations both upstream and within the coding of genes *Cii* and *Ciii*. Do these intra-coding sequences perhaps draw in EcRE-complexes to the gene region, thereby increasing their concentrations in the immediate region and so increasing responsiveness? Their presence within the coding of these two genes raises the intriguing possibility that their coding may effect more than the amino acid composition of their proteins.

The last exploration of the region’s sequence was to identify the boundaries of the event creating the inverted duplication of either gene *Cii* or *Ciii* within the region. A palindrome between *Cii* and *Ciii* in the center of the region (see Figure 1) seems likely to mark one end of the duplication, but might remnants of the *Ci* gene indicate its other end in the 5’ end of the region? Using the sequence CACCA that characterizes the *Ci* gene sequence, a search for its possible paleogenetic footprint in the region indicated multiple copies of the configuration within each of the genes and its rarity in intergenic sequence except for a cluster upstream of the *Ciii* gene site. Its presence confirms the possibility that some or all of the *Ci* sequence was within the originating duplication establishing genes *Cii* and *Ciii* in *99C*.

But more evidence of inclusion of *Ci* sequence in the duplication event within *99C* may be in the coincidence of the gene-like sequence in its 5’ region in the aligned searches of *99C*. Looking up through each of the juxtapositioned GCAT searches to the 2D dotplot of the region allows a visual collation of the CACCA, EcRE-like sites, G+C coding-like concentrations and finally the 2D checkerboard pattern that also characterizes the *Ci* gene site itself. This visual summation enabled by these alignments demonstrates another valuable facility of GCAT analysis. This is noted by the tentative designation of “*Cghost?”* on the map of the region’s annotations (Figure 1), as indication of some evidence of the birth and death of a once genetically active sequence.

These characterizations of the *Jonah 99C* region laid the ground for exploring two distinctively different yet possibly related kinds of genomic sequence and questions: introns, as found in the *Dip*-*B* gene, and transposable elements, using an apparently currently active element of the abundant transposable element (TE), *roo*, as examples. Each provided interesting, unexpected and perhaps functionally significant similarities and differences.

Their comparison arose from the curiosity of their high A+T-composition. As evident in their GCAT barcoding (Figures 2 and 3), *Dip*-*B’s* introns (A+T = 64.5%) and *roo*’s 7 kb ORF (A +T = 68.5%) are both topologically distinctively different from genomic coding (GCAT panels, Figures 1 and 2). This similarity led to exploring whether alternative excision might take place within these sequences. Might *Dip*-*B*’s large 5’exonic intron, for example, be alternatively excised in response to environmental signals, to provide alternative coding and activity, and how might an AT-dense ORF of 7 kb provide its coding?

Searches for the sequences associated with intron excision were therefore pursued, and as shown in Figures 2 and 3, the results were interestingly different. In *Dip*-*B’s* long intron, a second complement of the common TTAAT configuration, followed successively by a polypyrimidine tract and an AG excision site positioned to re-establish the coding of the originating ATG of the ORF were found indeed to be present near the annotated 3’ end of the long intron. But its consequent additional coding would include an IN-frame STOP codon! Thus, should alternative excision take place, its transcription would fail, saving the fly from the likely encumbrance of “mis-coded” alternative dud protein.

The same search of *roo’s* ORF (Figure 3) provided a different interesting result. TTAAT and its associated sequence configurations defining transcript excision are indeed found within the ORF at 7 sites. Further, several of these sites could allow independent excision of transcript sequence coding *roo*’s reverse transcriptase (RvT) and integrase (integ) (see Figure 3). Some corresponding evidence of their excision may be in the fragmentary *roo* transcript sequences found among the RNAs sampled in early embryos as indicated in the annotation of *roo{419}* (www,flybase.org). These coincident observations do confirm active transcription of the remarkable *roo* ORF, and further suggest that within its long ORF it carries “mini-ORFs” that may enable independent translation of parts of its coding.

The distribution of T and A in STOP codons in *Dip*-*B’s* introns and *roo*’s long ORF are differently notable. The long intron of *Dip*-*B* is packed with stop codons (TAA, TAG and TGA), many of which are IN-frame with the exon’s originating ATG. Thus their positioning stops translation if and/or when intron excision fails. In contrast, the 7 kb ORF of *roo* is also packed with stop codons, but each of these hundreds distributed throughout the ORF is out-of-frame with its apparently originating ATG. While the positioning of these striking AT-dense sequences may bring different functional consequences in the two sequences, the simple detail of their similar nucleotide composition, as made conspicuous in their GCAT profiles, is worth further consideration.

Common to both *Dip*-*B’s* introns and the long ORF of *roo* is their capacity to carry nucleotide that does not apparently provide genetic contribution to fly or TE. But do they provide a more basic metabolic function? *Dip*-*B’s* introns include about one third of the transcript’s nucleotide only to be discarded in the maturation of its mRNA. Within transcribed introns these nucleotides are osmotically-safe for the cell yet accessible for uptake in further transcription. This dynamic storage of nucleotide might well provide a strategic metabolic role in enabling transcription. The long ORF of *roo* might also provide a similar role. In this its abundance of out-of-frame STOP codons may be coincidental to its nucleotide composition as also a dynamic repository of nucleotide to be released from its transcript with excision of its translatable coding (see abve) as for further transcription.

This possibility of both sequences functioning as dynamic reservoirs of nucleotide substrates makes functional sense of these sequences as constituent parts of the fly genome, and may indeed also suggest a common history in the fly. Invasions of TEs including juxtaposed configurations allowing their excision might, as their similar structures suggest, become permanent fixtures in genes together with their source of transcriptional substrate! In any case, these possibilities are presented here as example of how visualization of sequence relationships often prompt valuable speculation.

Finally, it is hoped that these various examples of GCAT representation of sequence enabling visual identification of sequence configurations demonstrate their value as a complement to the more structured computational analyses of how this remarkable molecule of DNA provides its functionality.

While here these analyses have been carried out using old programs on an old Mac system, it is hoped that their demonstrations of GCAT’s value may prompt updating programs for use on present systems. One program recently offered by Seibt, Schmidt and Heitkam (2018) that they describe as a “customizable, ambiguity-aware” program and named “Flexidot” may answer the call.

## Acknowledgements

We thank Mary-Lou Pardue and Richard Lewontin for their interest and encouragement in this work.

